# HERC1 regulates breast cancer cells migration and invasion

**DOI:** 10.1101/2020.11.05.369975

**Authors:** Fabiana A Rossi, Ezequiel H Calvo Roitberg, Juliana H Enriqué Steinberg, Molishree Joshi, Joaquín M Espinosa, Mario Rossi

**Author notes:** Corresponding author: Mario Rossi, Translational Medicine Research Institute (IIMT-CONICET), Austral University, Buenos Aires, Argentina. Av. Perón 1500, Pilar (zip code: B1629AHJ), Buenos Aires, Argentina.

## Abstract

Tumor cell migration and invasion into adjacent tissues is one of the hallmarks of cancer and the first step towards secondary tumors formation, which represents the leading cause of cancer-related deaths. This process is considered an unmet clinical need in the treatment of this disease, particularly in breast cancers characterized by high aggressiveness and metastatic potential. In order to identify and characterize genes with novel functions as regulators of tumor cell migration and invasion, we performed a genetic loss-of-function screen using a shRNA library directed against the Ubiquitin Proteasome System (UPS) in a highly invasive breast cancer derived cell line. Among the candidates, we validated *HERC1* as a gene regulating cell migration and invasion. Furthermore, using animal models, our results indicated that *HERC1* silencing affects primary tumor growth and lung colonization. Finally, we conducted an *in-silico* analysis using publicly available protein expression data and observed an inverse correlation between HERC1 expression levels and breast cancer patients’ overall survival.

Altogether, our findings demonstrate that HERC1 might represent a novel therapeutic target for the development or improvement of breast cancer treatment.

## INTRODUCTION

The development of metastasis represents the principal cause of death in patients with solid tumors (1). The invasive potential of malignant cells is characterized by a series of molecular and genetic modifications that increase their invasive potential, allowing them to penetrate the extracellular matrix and spread in the surrounding tissues (2). These changes are associated with alterations in a wide variety of posttranslational modification patterns, including ubiquitylation (3). The Ubiquitin Proteasome System (UPS) plays a fundamental role in almost every single cellular process, and therefore it is not surprising that genetic alterations, abnormal expression or dysfunction of different components of this cascade are often associated with malignant transformation of tumor cells and the development of metastatic processes (4–10).

In order to find novel therapeutic targets for the treatment of cancer, we focused our attention on the identification of genes related to the UPS that had novel functions as positive regulators of migration, using a triple negative breast cancer (TNBC) derived cell line. This subtype of breast cancer (BC) is characterized by high metastatic rates, hormonal independence and lack of targeted therapies (11–13).

We performed a shRNA genetic screen targeting different UPS genes, using a combination of Boyden chambers and *in vivo* experiments as functional assays. After the selection procedure we identified *HERC1* as a candidate positive regulator of tumor cell migration and invasion. *HERC1* is a ubiquitin ligase, and our *in vitro* and *in vivo* experiments demonstrated that its depletion decreases invasiveness in highly aggressive breast cancer cells. Moreover, an *in-silico* analysis of HERC1 protein levels in breast cancer patients’ samples indicated that high expression in primary tumors correlates with decreased survival, compared to low-expressing tumors. These results highlight the potential of *HERC1* as a novel putative therapeutic candidate for cancer treatment.

## RESULTS

### Phenotypic screening for positive regulators of migration

We implemented a loss-of-function screen following an experimental protocol using a TNBC-derived cell line (MDA-MB-231), and Boyden chambers and mice in order to identify positive cell migration regulators within the UPS (Fig. 1A). We generated stable cell lines infected with an shRNA library directed against over 400 UPS genes (using around 5 shRNAs per gene) or an empty vector as a control. We then subjected these populations of cells to *in vitro* migration assays and amplified the non-migratory cells. We repeated this experiment several times to further enrich our population of interest, using the non-migratory cells from each migration experiment for the subsequent cycle of selection (Fig. 1B). Once the proportion of non-migrating cells reached a *plateau*, we inoculated these poorly migratory cells in NOD/SCID mice through tail vein injection. After two months, we resected lungs and brains from injected mice, purified the human cells and analyzed the relative abundance of the shRNAs compared to the cells prior to injection. Using this approach, we identified four genes that were substantially less represented in the tissues-derived cell lines compared to the initially injected cells, thus indicating that their expression may be needed for *in vivo* invasion. Among the candidate genes, Appbp1, Usp26 and Fbxl20 had already been reported as positive regulators of metastasis (14–17), indicating that our strategy was successful. In contrast, the role of HERC1 in the control of tumor cell invasion has not been studied yet and therefore we decided to further investigate its potential as a novel molecular target in breast cancer.

**Figure 1.**
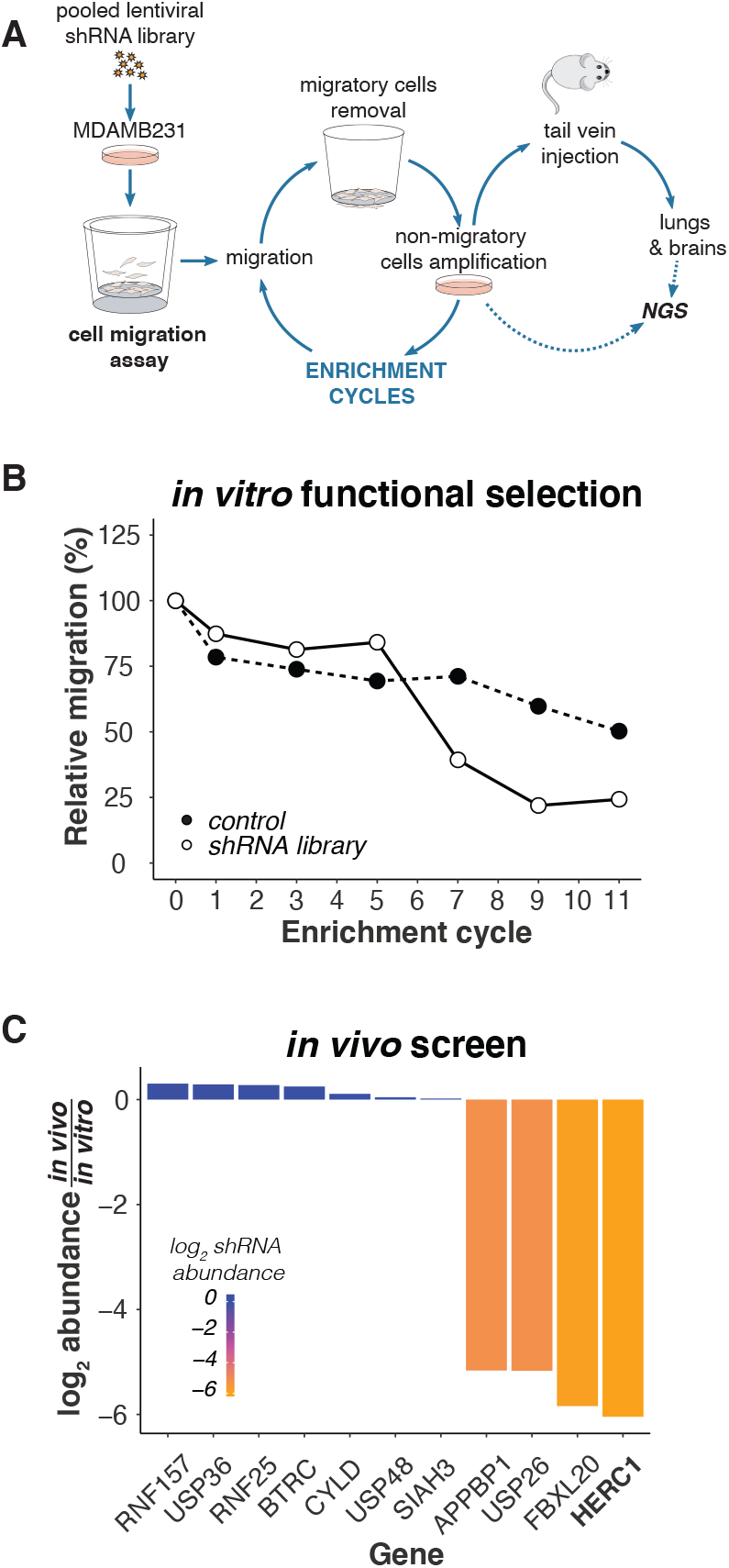
shRNA-based selection of positive regulators of cell migration. (**A**) Overview of the selection procedure. The production and infection of an ubiquitylation-related lentiviral shRNA library are described in Methods. Two weeks after lentiviral infection and selection, MDA-MB-231 cells were seeded onto Boyden chamber inserts and allowed to migrate across the porous membrane at 37°C for 24 hours in order to select cells with a decreased migration phenotype. Migrating cells were removed and non-migrating cells were collected by trypsin treatment from the inserts upper compartment and cultivated for one week. Cells were then reseeded onto Boyden chamber culture inserts for a subsequent round of selection; this procedure was repeated until cells lost 75% of their initial migratory potential. This non-migratory population of cells was then inoculated through tail vein injection into NOD/SCID mice, and primary cultures were generated after two months from lung and brain tissues. shRNAs were retrieved by PCR from the in vitro and in vitro/vivo-selected cells, and then identified by sequencing. (**B**) Boyden chamber assay was used every other enrichment cycle to determine the relative percentage of migratory cells and monitor the selection process. (**C**) shRNAs abundance after the in vivo selection, relative to their abundance in the in vitro-selected non-migratory cells.

### Proliferation is unaffected by *HERC1* silencing

Since alteration of proliferation rates might account partially for the selected phenotype, we measured doubling time of *HERC1* silenced and control cell lines. To this end, we generated *HERC1* knockdown (KD) cell lines with two different shRNAs (named shRNA#1 and #2) and used an empty vector-transduced cell line as a control (Fig. 2A). Our results indicated that *HERC1* KD does not affect proliferation in MDA-MB-231 cells (Fig. 2B). We further validated these results using another breast cancer cell line (Supp. Fig. 1).

**Figure 2.**
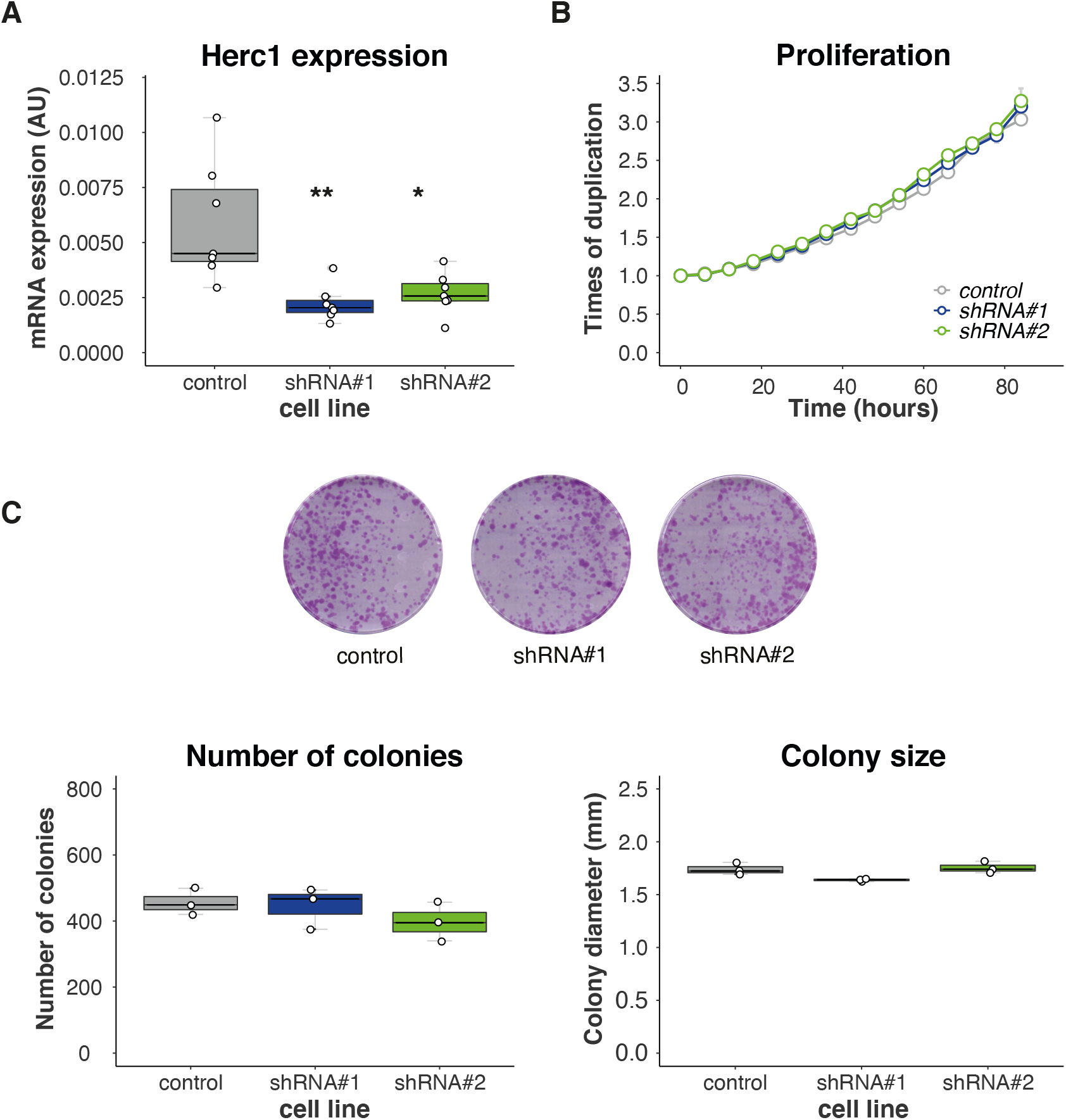
Proliferation is unaffected by HERC1 depletion. MDA-MB-231 cells were stably transduced with an empty vector (control) or two different shRNAs (#1 & #2) targeting HERC1. (**A**) Efficiencies of shRNA-mediated target gene knockdown were confirmed by RT-PCR (top, n= 7, one-way ANOVA, Dunnett’s multiple comparison test. shRNA#1 p= 0.0014 and shRNA#2 p= 0.0365). (**B**) An area-based microscopy method was used to determine cell growth over time. Cells were seeded onto wells and allowed to attach. At the indicated time points, cells were photographed, and the occupied area was calculated. The graph shows the occupied area relative to time= 0 hours. Doubling times= control: 47.01±0.2087, shRNA#1 46.23±0.3667, shRNA#2 47.38±1.023 (n= 3, Kruskal-Wallis, Dunn’s multiple comparison test. shRNA#1 p> 0.9999 and shRNA#2 p= 0.4503). (**C**) Cells were subjected to colony formation assay. Top: representative images of the crystal violet-stained 100 mm dishes at the end of the experiment. Bottom left: number of colonies generated after 15 days (n= 3, Kruskal-Wallis, Dunn’s multiple comparison test. shRNA#1 p> 0.9999 and shRNA#2 p= 0.4661), and bottom right: diameter of the resulting colonies (n= 3, Kruskal-Wallis, Dunn’s multiple comparison test. shRNA#1 p= 0.1473 and shRNA#2 p> 0.9999).

Next, we analyzed whether *HERC1* KD affects breast cancer cell clonogenicity. For that purpose, we plated cells at low confluency, and analyzed the resulting colonies after two weeks. We did not observe significant differences in neither colony number nor size between KD and control cell lines (Fig. 2C).

Altogether our results indicate that proliferation is unaffected by *HERC1* expression knockdown in TNBC cell lines.

### Validation of *HERC1* as a candidate gene regulating migration

In order to validate *HERC1* as a positive regulator of migration, we performed two independent cell migration assays.

Firstly, we subjected cells to Boyden chamber assay and observed a reduction in the number of migrating cells in *HERC1* KD lines, compared to the control cell line (Fig. 3A).

**Figure 3.**
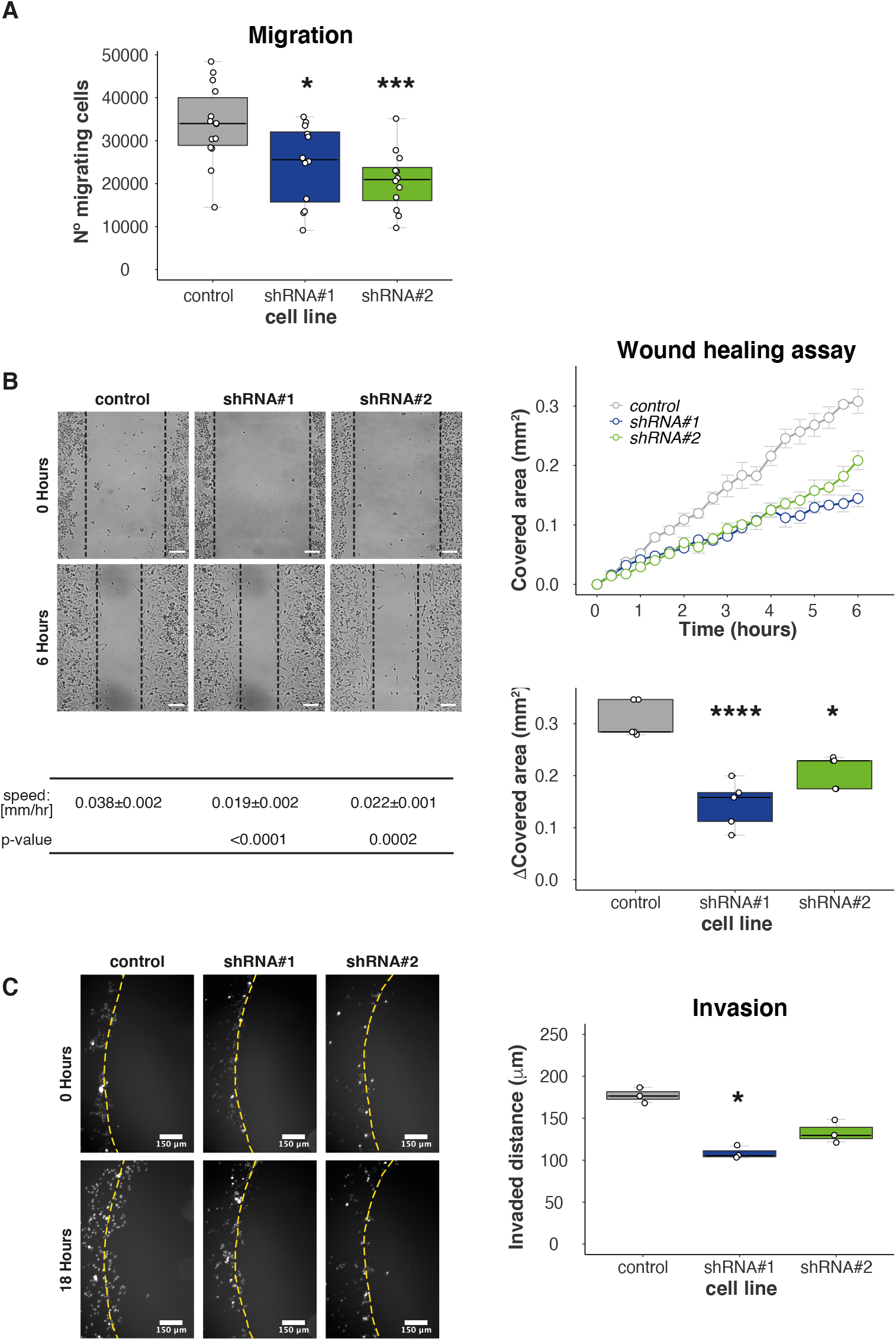
Validation and characterization of HERC1 as a candidate migration and invasion regulatory gene. Cells migratory potential was evaluated by two different experiments. (**A**) Boyden chamber assay: number of migratory cells per Boyden chamber membrane (n≥ 12, one-way ANOVA, Dunnett’s multiple comparison test. shRNA#1 p= 0.0180 and shRNA#2 p= 0.0009). (**B**) Wound healing assay. Top left: Representative areas in a wound healing experiment at the indicated time points, scale bar= 100 μm. Top right: wound covered area (mm^2^) at the indicated time points. Bottom left: wound edge closing speed (n= 5, one-way ANOVA, Dunnett’s multiple comparison test. shRNA#1 p< 0.0001 and shRNA#2 p= 0.0002). Bottom right: wound covered area (mm^2^) at endpoint (n= 5, one-way ANOVA, Dunnett’s multiple comparison test. shRNA#1 p< 0.0001 and shRNA#2 p= 0.0024). (**C**) Agar spot assay: cells were seeded in wells with drops of solidified agar and invaded along the bottom surface under the agar. Pictures were taken along the edge and the displacement is the extent of invasion under agar from the spot edge to cells final position. Left: Representative area showing cell invasion into an agar spot at the indicate time points. Scale bar= 150 μm. Right: cells mean displacement at the end of the experiment (n= 3, Kruskal-Wallis, Dunn’s multiple comparison test. shRNA#1 p= 0.0146 and shRNA#2 p= 0.3594).

Secondly, we used wound healing assays. To this end, we scratched a confluent cell monolayer using a pipette-tip and analyzed cells movement across the gap. Our results showed that *HERC1* KD significantly decreased the migratory potential of cells relative to their control (Fig. 3B). We observed similar results when another breast cancer cell line was used (Supp. Fig. 1).

Overall, these experiments indicate that *HERC1* silencing affects cell migration *in vitro*.

### *HERC1* regulates cell invasion

Next, we assessed cell invasiveness using agar spot assay and analyzed the distance covered by the cells inside the agar spot. We observed a significant reduction in *HERC1* KD cells displacement, compared to the control cell line (Fig. 3C). In addition, we performed this experiment using another breast cancer cell line, obtaining similar results. These observations indicate that *HERC1* KD-dependent alteration in migration also results in a significant reduction in breast cancer cell invasive potential.

### *In vivo* validation

In order to further characterize *HERC1*-dependent control of cell invasion *in vivo*, we performed two different experiments using immunocompromised mice.

We first injected MDA-MB-231 control or *HERC1*-silenced cells subcutaneously in the mammary fat pad of female mice and monitored tumor growth every 2-3 days. Tumor growth curves analysis indicated that those generated from MDA-MB-231 *HERC1*-silenced cells were significantly less volumetric than the ones originated from control cells. Moreover, Kaplan-Meier analysis of Tumor-Free Survival indicated that *HERC1* silencing affected tumor development (Fig. 4A).

**Figure 4.**
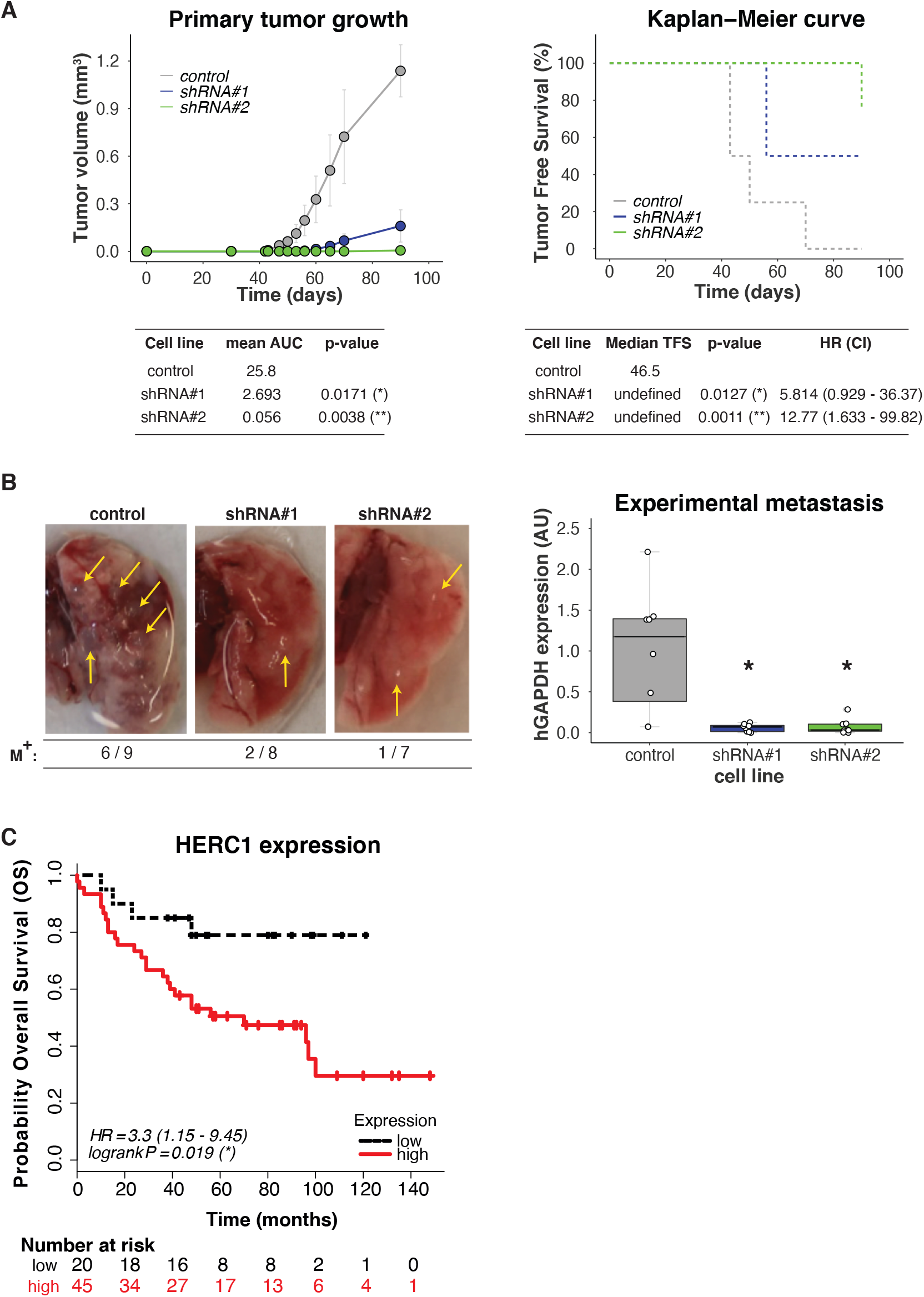
In vivo and in silico analysis of HERC1 relevance in tumor biology and patients’ survival. (**A**) Downregulation of HERC1 attenuates tumorigenicity in vivo. Control or HERC1-silenced MDA-MB-231 cells were subcutaneously inoculated into the mammary fat pads of female NOD/SCID mice and tumor growth was monitored every 2-3 days. Top left: tumor volume was calculated at the indicated time points (results show mean value ± s.e.m.), and bottom left: Area Under Curve (AUC) was performed in order to analyze differences between treatments (n≥ 4, Kruskal-Wallis, Dunn’s multiple comparison test. shRNA#1 p= 0.0171 and shRNA#2 p= 0.0038). Top right: Kaplan–Meier curves for Tumor Free Survival (TFS) in mice injected with control or HERC1-silenced cells, and bottom right: Long-Rank Test (Mantel-Cox) Analysis, HR: hazard ratio, CI: confidence interval (n≥ 4, Log-Rank (Mantel-Cox) test, shRNA#1 p= 0.0127 and shRNA#2 p= 0.0011). (**B**) Silencing effects of HERC1 on experimental metastasis assays: NOD/SCID male mice were inoculated with MDA-MB-231 control or HERC1-silenced cells through tail vein injection and lungs were harvested after 2 months. Left: representative images of lungs at endpoint; metastatic foci are indicated with arrows. M+: ratio of lungs positive for metastatic growth versus the number of injected mice. Right: metastatic potential was estimated by qPCR human DNA quantification, normalized to mouse DNA (n= 8, Kruskal-Wallis, Dunn’s multiple comparison test. shRNA#1 p= 0.0187 and shRNA#2 p= 0.0136). (**C**) Kaplan–Meier curves for breast carcinoma patients’ overall survival (OS) according to HERC1 protein expression status in primary tumors: red solid line and black dashed line indicate cases with high and low expression of HERC1, respectively (n=65, Log-Rank (Mantel-Cox) test, p= 0.019).

We then inoculated control or *HERC1*-silenced MDA-MB-231 cells through tail vein injection and harvested the lungs two months later. As shown in Figure 4B, *HERC1* depletion reduces metastatic load *in vivo,* as evaluated by human DNA quantification.

Our results indicate that *HERC1* silencing decreases cell engraftment and tumor growth, as well as colonization into the lungs.

### *In silico* analysis for HERC1 protein expression

Finally, we conducted an *in silico* analysis in order to study the association between *HERC1* protein expression levels in the primary tumors and breast cancer patient’s overall survival (OS), using an online KM-Plotter database. We built Kaplan-Meier plots by dividing the patients into two groups based on the expression of HERC1 protein, and observed a significant separation between the survival curves, with a worse prognosis associated with higher expression levels (Fig. 4C).

These findings indicate that HERC1 shows great potential as a new predictor of OS in breast cancer.

## DISCUSSION

Cell migration and invasion are critical steps in the metastatic dissemination of tumor cells, a process that constitutes the principal cause of death due to solid tumors (1). In fact, it has been reported that the regulation of transcriptional pathways related to migration, rather than proliferation, are strongly associated with breast cancer patient’s survival (18), highlighting the importance of its modulation in the development of new treatments for the clinical management of this disease.

For that purpose, we performed a genetic screen and a functional assay that involved both in vitro and in vivo experiments, in order to find genes related to the UPS with novel functions as regulators of cell motility. Using this approach, we identified *HERC1* as a putative positive regulator of migration in human breast cancer.

*HERC1* is a HECT E3 ligase which is restricted to the cytoplasm and Golgi/vesicular-like membrane compartments (19). Mutations of *HERC1* gene or its deregulation have been associated with different pathologies, including neurological disorders (20–31) and cancer (32, 33).

Our *in vitro* validation experiments showed that *HERC1* depletion did not affect cell proliferation but directly inhibited cellular migration in Boyden chamber and wound healing assays, as well as invasion. In the course of our studies, Herc1 was reported as a regulator of migration, further supporting our findings (34).

*In vivo* studies using immunocompromised mice demonstrated that *HERC1* silencing decreased tumor growth and colonization into the lungs in experimental metastasis assays.

Finally, we investigated the prognostic value of HERC1 protein expression in patients with breast cancer using the Kaplan-Meier plotter database. Our study indicated that higher levels of HERC1 protein are significantly associated with decreased overall survival, compared to patients that express low levels of HERC1. In concordance with our results, it has been demonstrated that HERC1 protein shows increased expression in tumor cell lines (19) and tumor breast cancer patients’ samples (35), compared to their respective normal counterpart.

Altogether our results indicate that *HERC1* has great potential as a putative new target for the treatment of breast cancer, and that its modulation could directly or indirectly regulate tumor cell motility and therefore patient’s outcome. Since HERC1 harbors catalytic activity, it would be attractive to further study whether its role in migration regulation is dependent on its E3 ligase activity, in order to better determine the most adequate strategy for its pharmacological inhibition. Lastly, our results highlight the importance of functional migration-based screens, as a complement to assays based on proliferation, for the discovery of putative new targets for the treatment of tumorigenesis.

## MATERIALS AND METHODS

### Cell lines and cell culture

The human breast cancer MDA-MB-231, MDA-MB-436 and embryonic kidney Hek293T cell lines were obtained from the ATCC and cultured in Dulbecco’s modified Eagle’s medium (DMEM) (Gibco) supplemented with 10% fetal bovine serum (FBS) (Natocor, Córdoba, Argentina), 50 U/ml penicillin-streptomycin and 200 μM L-glutamine at 37°C and 5% CO_2_ in a humidified incubator. ATCC uses morphology, karyotyping, and PCR based approaches to confirm the identity of human cell lines. Mycoplasm contamination was evaluated monthly by PCR, and cell lines were cultured less than three months.

### shRNA screening

A pool of plasmids encoding 1,885 shRNAs targeting 407 different components related to the ubiquitylation pathway inserted in the pLKO.1 backbone produced by The RNAi Consortium (TRC, Sigma-Aldrich, St. Louis, MO), was obtained from the University of Colorado Cancer Center Functional Genomics Shared Resource. 1 mg of the shRNA library plasmid DNA at 100 ng/mL was mixed with 4 mg of packaging plasmid mix (pD8.9 and pCMV-VSVG lentiviral packaging plasmids at a 1:1 ratio) and incubated with 30 mg of Polyethylenimine for 15 minutes at RT. The entire mixture was then added to a 100 mm dish containing Hek293T packaging cells at 75% confluence. 6 hours after transfection, media on cells was replaced with complete DMEM and 48 hours after media replacement, the supernatant from each dish of packaging cells (now containing lentiviral library particles) was filtered through 0.45 mm cellulose acetate filters and stored at −80°C until use. Target cells were seeded at 8×10^5^ cells/well in 6-well plates and then transduced with the lentivirus. After 48 hours the infective media was removed, and target cells were selected for 5 days with a 0.5 μg/ml puromycin DMEM medium. Cells were propagated for 14 days before use.

The following shRNAs were used for the generation of stable cell lines containing individual shRNAs targeting HERC1: TRC number 7243, sequence: 5’ GCCACATTTAGTTACTCTAAA 3’ (shRNA#1); TRC number 7244, sequence: 5’ GCTGATAAACTGAGTCCCAAA 3’ (shRNA#2).

### shRNA library preparation and sequencing

The library preparation strategy uses genomic DNA and two rounds of PCR in order to isolate the shRNA cassette and prepare a single strand of the hairpin for sequencing by means of an XhoI restriction digest in the stem loop region. After purity analysis of the sequencing library, barcode adaptors were linked to each sample to allow a multiplexing strategy. A HiSEQ 2500 HT Mode V4 Chemistry Illumina instrument was used for that purpose and each sample was quantified and mixed together at a final concentration of 10 ng/mL. Samples were sequenced with a simple 1×50 run and on average 1.2×10^6^ reads were obtained per sample (>600X shRNA library complexity).

### shRNA screen NGS data analysis

shRNA screen NGS data was analyzed in a similar fashion to RNA-seq data. Briefly, quality control was performed with FastQC, reads were trimmed to include only shRNA sequences using FASTQ trimmer and filtered with the FASTQ Quality Filter. Reads were then aligned to a custom reference library of shRNA sequences using Bowtie2.

### Boyden chamber migration assay

Cells were starved for 24 hours (DMEM 0.1% FBS). After trypsinization, 5×10^4^ were added to the top chamber of 24-well Boyden chambers (8 μm pore size membrane; BD Bioscience, Bedford, MA), and medium was added to the bottom chambers and incubated for 24 hours (DMEM 10% FBS). Non-migratory cells were removed, and following membrane fixation, cells were stained with 4ʹ,6-diamidino-2-phenylindole and mounted. The membrane was then fully imaged using an Axio Observer Z1 Inverted Epi-fluorescence microscope (Zeiss) with montage function. Image analysis was performed with Fiji software, using an automated analysis macro to measure the number of nuclei per Boyden chamber.

For the migration screen, 6.6×10^5^ starved MDA-MB-231 cells expressing the shRNA library (or control cell line) were plated onto 6-well Boyden chambers (8 μm pore size membrane; BD Bioscience, Bedford, MA) in assay medium. After 24 hours of incubation, the non-migratory cells were collected from the upper chamber, propagated and allowed to re-migrate eleven times for enrichment purposes. The non-migratory cells of 8 Boyden chambers were combined per each selection cycle (to ensure a >700 library coverage) and the percentage of non-migratory cells in each case was determined in 24-well plates as described before.

### quantitative PCR

Total RNA was extracted from cell lines using TRIzol reagent (Invitrogen) according to the manufacturer’s instructions and complementary cDNA synthesis was carried out using M-MLV reverse transcriptase in the presence of RNasin RNase inhibitor (Promega) and an oligo(dT) primer (Invitrogen).

Total DNA was extracted from cell lines, primary cultures or fixed tissues using DNeasy Blood and Tissue kit (Qiagen).

Quantitative real time PCR amplification, using specific primer sets at a final concentration of 300 nM, was carried out using the FastStart Essential DNA Green Master kit (Roche) at an annealing temperature of 60°C for 35 cycles, and a CFX96 PCR Detection System (Biorad). PCR primers were all intron spanning for mRNA expression analysis; expression was calculated for each gene by the comparative CT (∆CT) method with GAPDH for normalization.

Primer sequences are as follows. HERC1: 5’ AGGTTAAGCTGGTTGGAGAAG 3’ (Fw), 5’ GGCATTGGGAGAGGGTATAAG 3’ (Rv) and GAPDH: 5’ TGCACCACCAACTGCTTAGC 3’ (Fw), 5’ GGCATGGACTGTGGTCATGAG 3’ (Rv) for cDNA measurement. Human GAPDH: 5’ TACTAGCGGTTTTACGGGCG 3’ (Fw), 5’ TCGAACAGGAGGAGCAGAGAGCGA 3’(Rv) and mouse GAPDH: 5’ CCTGGCGATGGCTCGCACTT 3’ (Fw), 5’ ATGCCACCGACCCCGAGGAA 3’ (Rv) for DNA measurement.

### Area-based analysis of proliferation rate

This experiment was based on an area-based imaging assay reported previously (36). Briefly, 1×10^3^ cells were seeded in 96-well plates and incubated overnight to allow cell attachment. Images were then acquired under bright field illumination every 6 hours for 3 days using a 10X air objective and Zen Blue 2011 software (Zeiss) for image acquisition. Image analysis was performed with Fiji software, using an automated analysis macro to measure the occupied area by cells.

### 2D clonogenic assay

1×10^3^ cells were plated on 100 mm dishes and medium was replaced every 3 days for 15 days. At this point, dishes were washed with 1X PBS and methanol-fixed for 15 minutes. Dishes were then re-washed and stained with a 0.05% crystal violet solution for 1 hour. Exceeding staining solution was then removed by immersion into a tap water containing recipient, after which dishes were air-dried, photographed and analyzed with the OpenCFU software.

### Wound healing assay

2.5×10^5^ cells were seeded in 24-well culture plates per well. Confluent monolayers were starved during 24 hours in 0.1% FBS-supplemented DMEM medium and a single scratch wound was created using a micropipette tip. 1X PBS was used to wash and remove cell debris, and then cells were incubated with 0.3% FBS-supplemented DMEM medium to enable migration through the wounds. Images were acquired using an Axio Observer Z1 Inverted Epi-fluorescence microscope (Zeiss), equipped with an AxioCam HRm3digital CCD camera, a Stage Controller XY STEP SMC 2009 scanning stage and an Incubator XLmulti S1 and Heating Unit XL S1 for live imaging incubation. Images were obtained under bright field illumination every 20 minutes for 6 hours, using a 10X air objective and Zen Blue 2011 (Zeiss) software for image acquisition. Image analysis was performed with Fiji software, using an automated analysis macro to measure the occupied area by cells, and the results are presented as the wound covered area at the end of the experiment, relative to time= 0 hours.

### Noble agar invasion assay

The procedure was performed as described previously, with minor modifications (37–39). A 1% noble agar solution was heated until boiling, swirled to facilitate complete dissolution, and then taken off of the heat. When the temperature cooled to 50°C, 5 µL spots were pipetted onto 96 well cell culture plates and allowed to cool for 20 minutes at RT under the hood. 5×10^3^ cells were then plated in the presence of 10% FBS cell culture media supplemented with 1 μg/ml Hoeschst 33258 (ThermoFisher Scientific) and allowed to adhere for 1 hour. Fluorescent images of the edges of each spot were taken every 20 minutes during 18 hours on an Axio Observer Z1 (Zeiss) Fluorescence Microscope using a 10X magnification air objective, equipped with CCD AxioCam HRm3 digital Camera, and a XLmulti S1 (D) incubation unit plus a XL S1 (D) temperature module to maintain cell culture conditions at 37°C and 5% CO_2_. Acquisition was controlled with Zen Blue 2011 (Zeiss) Software.

The number and position of cells was determined using image analysis software, ImageJ/Fiji ‘Trackmate’ plug-in (40).

### *In vivo* screen and experimental metastasis model

NOD/SCID mice were originally purchased from Jackson Laboratories (Bar Harbor, ME, USA), and bred in our animal facility under a pathogen-free environment. For all experiments, 7/8-week-old mice were used in accordance with protocols approved by the Institutional Board on Animal Research and Care Committee (CICUAL, Experimental Protocol # 63, 22.nov.2016), Facultad de Ciencias Exactas y Naturales (School of Exact and Natural Sciences), University of Buenos Aires.

For the *in vivo* screen, 1×10^6^ cells containing the shRNA library were resuspended in 200 μL of sterile 1X PBS and injected in the lateral tail vein of male mice. At endpoint, mice were euthanized, and lungs and brains were resected for isolation of primary cultures. Approximately 100 mg of tissue was grossly chopped with a scalpel, and the resulting mixture was suspended in 500 μl of 0.075 mg/ml Collagenase Type IA (Sigma) and incubated at 37°C. After 30 minutes, dissolved material was centrifuged at 1000 rpm for 5 minutes and the supernatant was discarded. The pellet containing tumor cells was then resuspended in 2 ml of red blood cells lysis buffer (155 mM NH_4_Cl, 10 mM KHCO_3_, 0.1 mM EDTA; pH 7.3) and incubated at 4°C for 3 minutes. 5 volumes of assay medium were then added, and the centrifugation step was repeated. The tumor cell sediment was then resuspended in 3 ml of assay medium, seeded in cell culture dishes and incubated at 37°C and 5% CO_2_.

For the experimental metastasis assay, 1×10^6^ HERC1 silenced or control cells were resuspended in 200 μL of sterile 1X PBS and injected in the lateral tail vein of male mice. 60 days post injection lungs were harvested, fixed in buffered formalin and stored in 70% ethanol until DNA extraction and metastatic load quantification following experimental procedures as described previously (41).

### Tumorigenesis model

For *in vivo* mouse tumor studies, 5×10^5^ transduced cells were suspended in 100 μl of sterile 1X PBS and subcutaneously injected in the mammary fat pads of female mice. Tumors were measured every 3 days and tumor volumes were calculated using the following formula: Vol (volume)= ½ (width^2^ * length). Area Under Curve (AUC) analysis was performed using measurements from mice that were alive at the end of the experiment.

### *In-silico* analysis

The KM-Plotter database contains proteomic data regarding the survival of 65 patients with breast cancer (Overall Survival (OS) data). The association between the HERC1 protein expression levels and OS was analyzed using an online KM-Plotter database(42) using the protein expression data and the survival information of patients with breast cancer downloaded from the GEO (43). Cohorts of patients were split by auto-select best cutoff expression values. All subtypes of breast cancer samples were included in the analysis.

The Kaplan-Meier survival plots with number at risk, hazard ratio (HR), 95% confidence intervals (CI) and log-rank p-values were obtained using the Kaplan-Meier plotter website (https://kmplot.com/analysis/index.php?p=service&cancer=breast_protein), accessed on October 28^th^, 2020.

### Statistical analysis

Results are presented as Box-and-whisker plots with median interquartile ranges plus minimum to maximum. *n* indicates the number of independent biological replicates. The one-way ANOVA with Dunnett’s multiple-comparison test as well as non-parametric Kruskal-Wallis and Dunn’s Tests were used to compare treatments to their corresponding control, and adjusted p-values are indicated. P-value differences of < 0.05(*), < 0.01(**), < 0.001(***) or < 0.0001(****) were considered statistically significant. GraphPad Prism statistical software (version 8.2.1) was used for the analysis.

## COMPETING INTERESTS

The authors declare no potential conflicts of interest.

## SUPPLEMENTARY FIGURE

**Figure S1.**
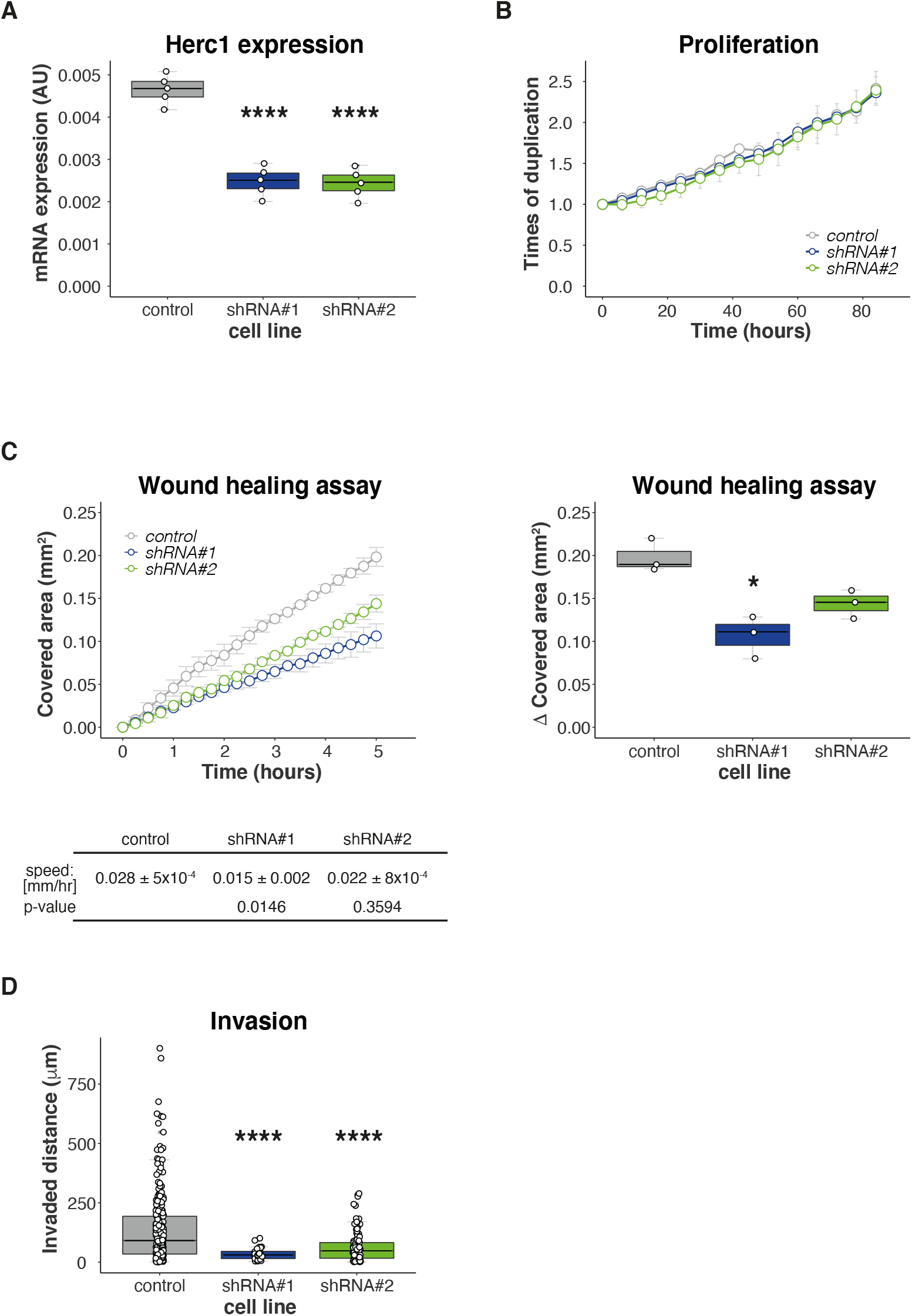
In vitro validation of HERC1 using MDA-MB-436 cells. (**A**) Efficiencies of shRNA-mediated HERC1 knockdown were confirmed by RT-PCR (top, n=5, one-way ANOVA, Dunnett’s multiple comparison test. shRNA#1 p< 0.0001 and shRNA#2 p< 0.0001). (**B**) An area-based microscopy method was used to determine cell growth over time. Cells were seeded onto wells and allowed to attach. At the indicated time points, cells were photographed. The graph shows the occupied area relative to time= 0 hours (Doubling times= control: 67.91±1.485, shRNA#1 62.99±1.508, shRNA#2 75.25±8.893. n= 3, Kruskal-Wallis, Dunn’s multiple comparison test. shRNA#1 p= 0.5480 and shRNA#2 p= 0.2485). (**C**) Wound healing assay was performed to analyze HERC1 silencing effects on MDA-MB-436 cells migration. Top left: wound covered area (mm^2^) at the indicated timepoints. (n= 3, Extra sum-of-squares F Test. shRNA#1 p< 0.0001 and shRNA#2 p< 0.0001), bottom left: wound edge closing speed (n= 3, Kruskal-Wallis, Dunn’s multiple comparison test. shRNA#1 p= 0.0146 and shRNA#2 p= 0.3594). Right: wound covered area (mm^2^) at endpoint (n= 3, Kruskal-Wallis, Dunn’s multiple comparison test. shRNA#1 p= 0.0225 and shRNA#2 p= 0.2721). (**D**) Agar spot assay was used to analyze invasion; the graph shows the cells displacement at the end of the experiment (n=1, more than 25 individual cells were measured per cell line. Kruskal-Wallis, Dunn’s multiple comparison test. shRNA#1 p< 0.0001 and shRNA#2 p< 0.0001).

## Notes

All authors have seen and approved the manuscript, and it hasn’t been accepted or published elsewhere. The authors declare no potential conflicts of interest.

### Competing Interest Statement

The authors have declared no competing interest.

